# Population-aware computational assessment of base-editing accessibility among pathogenic variants on human chromosome 21: an African-ancestry-focused pilot study

**DOI:** 10.64898/2026.11.12.766091

**Authors:** Ernest Achab-Awelana, Emmanuel Kodom Yeboah, Christian Tetteh Aryeh, Abednego Essandoh

## Abstract

**Background:** Population-aware gene-editing research requires independent evaluation of clinical variant evidence, ancestry-stratified allele observations, nucleotide correction chemistry, and sequence-level editing constraints. We developed a source-traceable chromosome 21 pilot connecting these dimensions without equating computational accessibility with therapeutic editability.

**Methods:** Reviewed ClinVar pathogenic/likely pathogenic single-nucleotide variants were matched by GRCh38 chromosome, position, REF and ALT to overall and African/African-American (AFR) gnomAD v4.1 exome allele counts. Exploratory frequency contrasts used two-sided Fisher exact tests and Benjamini–Hochberg correction. Higher-confidence variants with canonical adenine or cytosine base-editing chemistry underwent NGG PAM/editor-window screening, bystander enumeration, MANE v1.5 consequence annotation, theoretical joint-product assessment and chromosome 21–restricted guide-homology searches.

**Results:** Among 1,944 ClinVar variants, 851 matched gnomAD; 694 were quality-filtered and alternate-allele-observed overall, including 300 higher-confidence records. Six variants met exploratory AFR-elevation criteria. Of 234 higher-confidence ABE/CBE chemistry-compatible variants, 118 (50.4%) had at least one nominal NGG/PAM-window placement, 55 (23.5%) a nominal zero-bystander placement, and 44 (18.8%) a zero-bystander placement without an additional chromosome 21 NGG near-match within three mismatches. The 261 placements generated 295 hypothetical bystander events (195 distinct alleles); five distinct potential products overlapped pathogenic/likely pathogenic ClinVar alleles. Across placements, 764 theoretical target-corrected sequence states were enumerated, of which 503 contained at least one bystander. Three variant-level editing-accessibility comparisons between AFR-observed and AFR-unobserved alleles were not statistically significant (BH q=1.0).

**Conclusions:** Canonical correction chemistry substantially overestimates nominal local base-editor accessibility. A population-aware, transcript-informed computational framework can identify hypotheses for follow-up, but allele-frequency contrasts, guide-similarity results and enumerated sequence products do not establish clinical efficacy, experimental editing specificity or genome-wide off-target safety.

## Introduction

Genome editing has expanded from nuclease-mediated disruption to precision strategies capable of changing single nucleotides without requiring the same double-strand break-and-donor template paradigm. Cytosine base editors (CBEs), adenine base editors (ABEs), and prime editors differ in their chemical capabilities, target constraints and unintended-product profiles [1–4]. In clinical translation, however, a disease-associated substitution is not a viable editing target merely because its reference-restoration change belongs to an ABE-or CBE-compatible transition class. Sequence context, the protospacer-adjacent motif (PAM), activity window, bystander bases, the disease mechanism, delivery and safety must all be considered [3,10,11].

Large clinical and population databases provide complementary—not interchangeable— sources of information. ClinVar aggregates submitted clinical assertions, including their review status, whereas gnomAD reports observed variation and site-specific allele counts across sequencing cohorts [5–7]. The presence of an allele in a reference population does not establish its penetrance or disease burden; conversely, its absence may simply reflect limited coverage or sampling. Clinical interpretation also depends on condition-specific evidence, inheritance mechanism and the relationship between pathogenicity and the particular intended edit [7,18].

African populations encompass exceptionally broad genetic diversity, yet have historically been underrepresented in biomedical genomics [8,9]. The gnomAD AFR label is an inferred, heterogeneous genetic-ancestry grouping (African/African-American), not a census of residents of Africa, an ethnic identity, or an estimate of continent-wide disease prevalence [6,19]. Accordingly, a population-aware workflow should retain the actual allele counts and denominators, quantify uncertainty, and distinguish a signal that is relatively elevated in a sampled group from one that is clinically important.

Existing in-silico analyses often summarize guide availability without explicitly joining clinical confidence tiers, ancestry-specific observations, joint editing outcomes and transcript-level consequences. The biochemical targeting constraints and the possibility of correlated editing at adjacent bases make single-base flags an incomplete description of candidate editing products [11,12]. Furthermore, off-target sequence-similarity counts are sensitive to the scope of the reference genome and mismatch search; the absence of nearby homology on one chromosome cannot be generalized to genome-wide specificity [14].

We therefore developed a staged, source-traceable chromosome 21 pilot. Our primary objectives were to (i) characterize reviewed pathogenic/likely pathogenic single-nucleotide variants; (ii) compare observed AFR and pooled non-AFR exome allele frequencies; (iii) assess reference-restoration chemistry and nominal NGG-dependent base-editor accessibility; (iv) catalog putative bystanders, MANE transcript consequences and combinations of on-target changes; and (v) evaluate chromosome 21 guide homology under a defined mismatch threshold. We hypothesized that adding sequence and consequence constraints would substantially reduce the subset passing simple chemical classification, while treating ancestry-related differences in accessibility as an empirical question rather than a presumption.

The translational rationale for such screening has sharpened as genome editing moves beyond proof of concept. Genome-edited autologous hematopoietic stem-cell therapies have provided clinical evidence that targeted genetic modifications can confer meaningful benefit in severe hemoglobinopathies, and a personalized in-vivo base-editing treatment was recently administered to a child with carbamoyl phosphate synthetase 1 deficiency. A 2026 early clinical report of HBG promoter base editing in sickle-cell disease further illustrates the pace of platform development. These achievements should not, however, be conflated with therapeutic feasibility at an unrelated locus: editing modality, delivery, tissue turnover, required allelic correction fraction, and the clinical architecture of the disorder differ substantially [28–30,39,40].

A precise distinction must also be maintained between “base-editor-compatible chemistry” and “correctable disease genotype”. The A→G or C→T reaction is a molecular capability of an editor; successful variant correction requires an appropriate allele, an accessible PAM, productive positioning inside a context-dependent activity window, acceptable collateral products and, ultimately, a biologically sufficient rescue. Conversely, a variant excluded by a canonical NGG screen is not necessarily refractory to all contemporary CRISPR-derived tools. PAM-relaxed Cas9 variants, engineered editors and prime-editing architectures enlarge the sequence space but introduce different activity and specificity trade-offs [22,23,38,41].

The population dimension presents a second interpretive challenge. Genetic diversity is not distributed uniformly across or within continents; African genetic variation is particularly extensive and incompletely sampled in widely used databases. Genome-wide studies across African ethnolinguistic groups show that additional sequencing uncovers substantial previously undescribed variation, and studies of global diversity emphasize how heterogeneous populations can be obscured by broad ancestry labels [24–27]. Thus, a variant’s relative frequency in gnomAD’s genetically inferred AFR group is a hypothesis-generating description of sampled alleles, not a direct estimate of disease burden, local prevalence or the probability that a patient in a specific African community carries the allele.

There are additional reasons to resist a simplistic “editable or not editable” dichotomy. Base-editing reaction outcomes are highly dependent on the arrangement of deaminatable nucleotides, editor configuration and the edited product that emerges. High-throughput outcome studies demonstrate that single-locus sequence context can substantially influence product mixtures, making the mere presence of a bystander base an imperfect surrogate for editing purity [11,12]. Moreover, a predicted synonymous bystander may reside close to a splice junction, while two changes in the same codon can generate a combined amino-acid outcome that cannot be inferred by independently annotating each substitution [13,34].

Accordingly, we structured the present study as an explicitly nested computational evidence hierarchy. Clinical review status is separated from population observation; population observation from nucleotide chemistry; chemistry from local PAM-window accessibility; local accessibility from theoretical product space; and local sequence homology from biological off-target activity. This hierarchy allows informative negative findings and failed screening gates to be reported rather than hidden, and it provides a transparent basis for targeted follow-up research without treating computational selection as a claim of experimental therapeutic readiness.

## Materials and methods

### Study design, source releases and reproducibility

This retrospective computational pilot used ClinVar GRCh38 VCF records with source date 1 November 2026; matched gnomAD v4.1 chromosome 21 exome overall and AFR allele-count records; the authentic UCSC GRCh38/hg38 chromosome 21 reference sequence (46,709,983 bp); and MANE GRCh38 version 1.5 transcript annotation. Analyses were restricted to 1-based chromosome 21 A/C/G/T biallelic SNVs, and no human participant-level phenotypes, interventions or experimental editing measurements were available. The source files, derived outputs, software and quality-control checksums were maintained as a versioned project archive.

Every variant was identified by (chromosome, 1-based position, reference allele, alternate allele). We did not lift over between genome assemblies, infer population frequencies for unmatched sites, or treat a GRCh38 reference allele as necessarily function-restoring in the absence of mechanistic evidence.

### ClinVar filtering and clinical-confidence stratification

We selected biallelic, canonical A/C/G/T SNVs with aggregate germline significance exactly “Pathogenic”, “Likely pathogenic” or “Pathogenic/Likely pathogenic” and ClinVar review status of at least one star. The prespecified primary evidence tier comprised 2–4-star records (multiple submitters without conflict, expert panel, or practice guideline); one-star records were retained for expanded/sensitivity analyses. Aggregate classifications were not treated as a substitute for condition-specific RCV/SCV assertions [5,18]. The clinical selection was not population-sampled.

### gnomAD allele joining, quality control and genetic-ancestry comparison

ClinVar alleles were exactly joined to the gnomAD v4.1 exome VCF using GRCh38 chromosome, position, REF and ALT. The primary frequency analyses required a PASS site filter and positive overall alternate-allele count. For matched sites, non-AFR remainder counts were derived as AC_remainder = AC_total − AC_AFR and AN_remainder = AN_total − AN_AFR, after verifying nonnegative counts and denominators. Frequencies were AC/AN, and relative frequency was AF_AFR/AF_remainder when calculable. “Remainder” contains all non-AFR-inferred exome contributors; it is not specifically a European or East Asian cohort.

Two-sided Fisher exact tests were applied to allele-count 2×2 tables and p values adjusted by the Benjamini–Hochberg procedure within declared test families (694 complete evaluable records; 300 higher-confidence records). Exploratory AFR-elevation flags required AFR AC ≥5, AFR/remainder frequency ratio ≥2 and q <0.05. Threshold sensitivity examined AFR AC cutoffs of 3, 5 and 10. Exact Clopper–Pearson 95% intervals were calculated for selected allele-frequency estimates. Analyses are susceptible to cohort and ancestry-structure confounding; q-values were not interpreted as causal evidence [6,7,15–17].

### Reference-restoration chemistry and sequence-context screening

The proposed correction was defined conceptually as ALT→REF, with A→G or T→C categorized as canonical ABE-compatible and C→T or G→A as canonical CBE-compatible. Other SNV corrections were categorized as noncanonical for these two systems, without presuming that prime editing or another approach would necessarily succeed. For each chemistry-compatible, 2–4-star variant observed in overall gnomAD, the GRCh38 reference base was checked against the uploaded FASTA and the corresponding single-ALT local sequence was reconstructed.

We enumerated 20-nt protospacers next to canonical SpCas9 NGG PAMs in both orientations, retaining placements with the target nucleotide in nominal windows: ABE7.10 positions 4–7, ABE8e positions 3–11 and BE4 positions 4–8, measured from the PAM-distal end. These are deliberately simplified screening windows, not fitted per-sequence activity models. Other editable adenines or cytosines in the same model window were counted as nominal bystanders; a zero-bystander option meant no other same-class editable base inside that *specified* window [1–3,10–12].

### Transcript-aware annotation and clinically flagged bystanders

Potential bystander substitutions were represented as plus-strand GRCh38 REF→ALT alleles, verified against the authentic chromosome 21 reference, and deduplicated by genomic variant key. Genomic intervals were intersected with MANE GRCh38 v1.5 transcript annotations to classify coding, synonymous, missense, UTR, intronic and splice-proximal outcomes. Coding effects were translated on the annotated transcript strand, with compound substitutions within the same codon evaluated jointly. Exact-coordinate/allele overlaps with the study ClinVar pathogenic/likely pathogenic records were recorded as clinical-review flags, not as evidence that the hypothetical products would occur or cause disease.

### Chromosome 21 homology and integrated filtering

Unique 20-nt guide sequences were compared with every A/C/G/T-only NGG-associated protospacer on both strands of GRCh38 chromosome 21, allowing zero through three ungapped Hamming mismatches. The intended genomic site was excluded from additional-site counts, and a reference-limited low-homology screen was combined with zero nominal bystander status *at the same guide placement*. The streaming implementation was independently regression-tested against the original chromosome 21 scanner. This is a site-similarity census—not predicted off-target editing—and excluded other chromosomes, non-NGG PAMs, bulges, genomic polymorphisms and chromatin [14].

Variant-level association tests compared AFR-observed (AFR AC>0) with AFR-unobserved (AFR AC=0 in the sampled subset) among the 234 primary chemistry-compatible variants for (i) any nominal PAM/window placement, (ii) any zero-window-bystander placement and (iii) an option satisfying both the zero-bystander and the chromosome 21–restricted no-additional-near-match criteria. Two-sided Fisher exact tests were reported with conditional odds-ratio estimates and exact conditional 95% confidence intervals; p values from the three comparisons were BH adjusted. These were exploratory comparisons of selected variants, not tests of equivalence or population-level editing efficiency.

### Allele-number uncertainty, homozygote counts and threshold sensitivity

For each quality-filtered matched locus, we retained gnomAD overall and AFR alternate-allele counts (AC) and allele numbers (AN), checked that AFR counts did not exceed overall counts, and calculated remainder counts by subtraction only when the cohort definitions were compatible. Clopper–Pearson exact 95% binomial confidence intervals were generated for group allele frequencies; as these are descriptive aggregate allele-count intervals, they do not correct for relatedness or population sampling structure. The number of observed alternate-allele homozygotes was also audited by cohort where genotype counts were explicitly available. The absence of observed homozygotes was not interpreted as absence of homozygous people in the underlying populations [7,16,24].

Because extreme relative frequencies can arise from sparse alternate-allele observations, AFR enrichment flags were interrogated at alternative minimum AFR allele counts of ≥3, ≥5 and ≥10 while retaining the predefined ratio threshold and FDR criterion. This sensitivity analysis was not intended to select a single biologically privileged cutoff; rather, it estimates the dependence of the descriptive candidate list on an arbitrary observation threshold. Specific disease frequency, inheritance and clinical ascertainment were not available for case-level interpretation.

### Joint editing-state enumeration and nonadditive codon outcomes

For each nominal editor-specific placement with b additional editable bases in the assumed window, all 2^b bystander-subset states were enumerated conditional on intended correction; the same subsets were also enumerated with the original target allele retained, giving 2×2^b theoretical states per placement. Each state was represented on the simplified reference-background local sequence with the ClinVar ALT target allele, and simultaneous substitutions within a coding triplet were translated together. This is a finite combinatorial census—not an empirical or machine-learning estimate of the probability of a particular molecular product [11,12].

Outcome annotation flags were non-mutually exclusive: a state could contain multiple nonsynonymous changes, an essential splice-junction substitution, and an exact ClinVar-overlapping allele. Counts were therefore stratified by the analysis unit and never summed as if they represented distinct patients, unique pathogenic variants or expected editing events.

### Reproducibility, regression checks and evidence boundaries

Independent implementations verified source REF alleles, coding-phase consistency, protein translation and chromosome 21 NGG near-match enumeration. The homology program used a four-block seed-and-verify strategy to guarantee retrieval of 20-nt protospacer matches within at most three ungapped substitutions under the specified NGG-only model. A memory-bounded streaming implementation reproduced all 256 intended and additional guide–site matches from an independently generated chromosome 21 audit. These technical checks establish internal computational reproducibility, not external biological validation [14,35,36].

We reported descriptive counts, Fisher exact p values and Benjamini–Hochberg q values without assigning causal interpretation to observational frequency differences. We did not impute unobserved AFR allele frequencies, infer allele phase, assume pathogenicity for all possible edited outcomes or run a whole-genome off-target search. Variants with insufficient sequence or external annotation evidence retained an explicit unevaluated status rather than being coded as unfavorable or safe.

## Results

### Cohort definition and matching

The ClinVar selection yielded 1,944 reviewed pathogenic/likely pathogenic chromosome 21 SNVs, comprising 732 in the 2–4-star primary tier and 1,212 one-star records. ABE-compatible reference-restoration chemistry was assigned to 865, CBE-compatible chemistry to 245, and other transitions/transversions to 834; thus 1,110/1,944 (57.1%) met chemistry-only ABE/CBE criteria. Exactly 851/1,944 matched gnomAD exome alleles; 694 were PASS with overall AC>0, of which 300 were higher-confidence. The intersection of the 300 primary variants with canonical chemistry yielded 234 candidates (194 ABE; 40 CBE). These sequential denominators refer to different eligibility conditions and must not be conflated. The analysis sets are detailed in Table 1.

**Table 1.** Definition of analysis sets and computational screening outcomes.

| Set / criterion | Variants, n | Interpretation |
|---|---|---|
| ClinVar reviewed P/LP SNVs (≥1 star) | 1,944 | Initial GRCh38 chr21 clinical set |
| Higher-confidence ClinVar (2–4 stars) | 732 | Prespecified primary clinical evidence |
| Exact overall gnomAD v4.1 matches | 851 | Exact GRCh38 allele join |
| PASS and AC_total >0 | 694 | Paired population-frequency analysis |
| Higher-confidence among 694 | 300 | Primary observed set |
| Set / criterion | Variants, n | Interpretation |
| ABE/CBE chemistry among 300 | 234 | Sequence screening denominator |
| NGG PAM + nominal editor window | 118 | At least one editor-specific placement |
| At least one nominal zero-bystander option | 55 | Window-limited local sequence descriptor |
| Same-guide zero-bystander + chr21 low-homology | 44 | Not genome-wide safety qualified |

Note. P/LP = pathogenic/likely pathogenic; all quantities count distinct variants, not editing placements.

Progressive restriction of eligible variants is illustrated in Fig 1.

**Fig 1.**
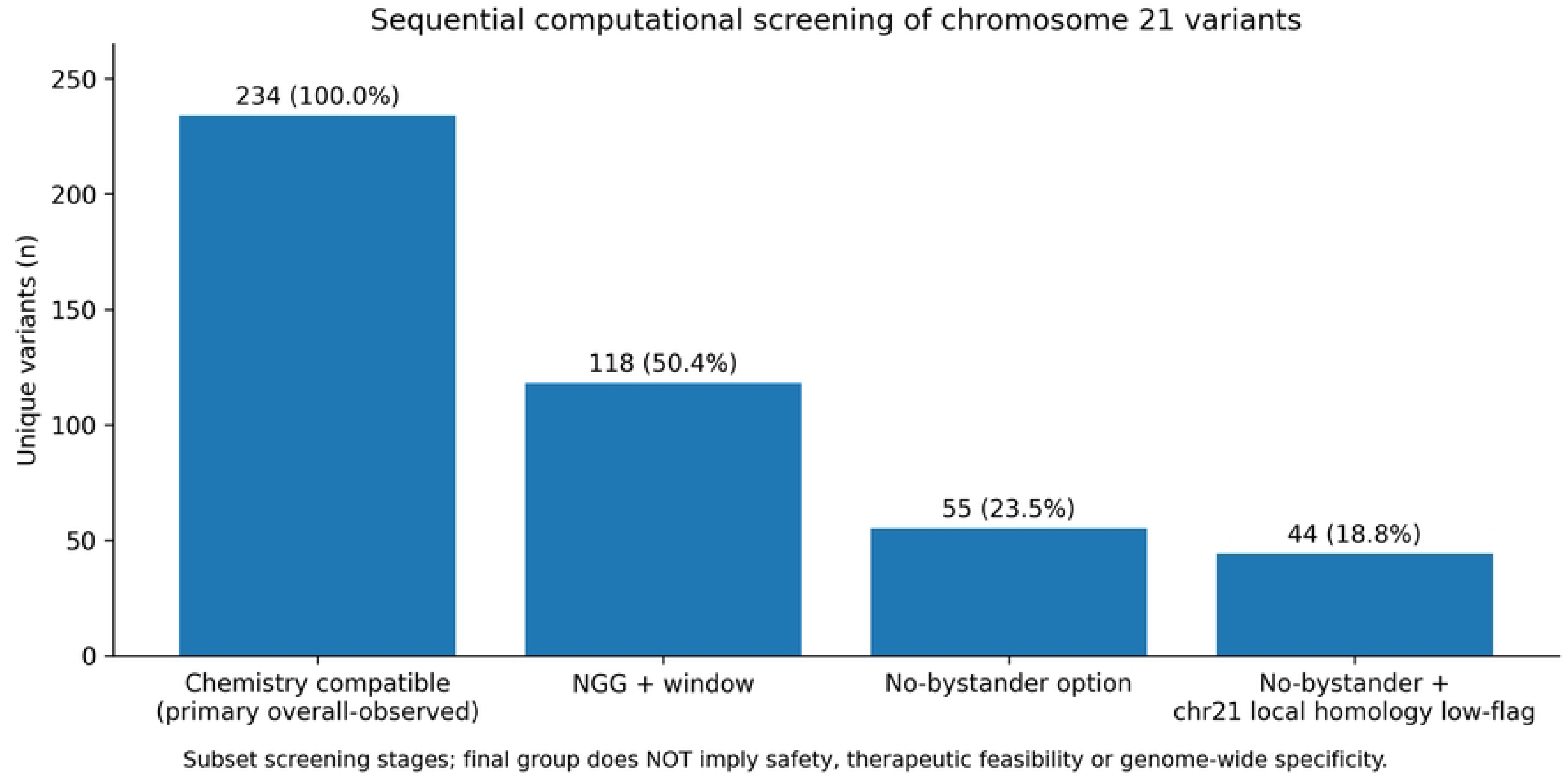
Sequential filtering of chemically compatible chromosome 21 variants. Sequential attrition of the 234 higher-confidence, overall-gnomAD-observed, canonical ABE/CBE-chemistry-compatible variants as increasingly restrictive sequence-level filters were applied. The final stratum is chromosome 21-limited and not a safety prediction.

### Genetic-ancestry allele frequencies and exploratory elevated variants

Among 694 evaluable overall-observed sites, 109 had a positive AFR allele count. Twenty-six allele-count contrasts met BH q<0.05 across the 694-test family (in either direction); six additionally met the explicitly exploratory AFR-elevation criteria. Four of these six belonged to the 300-variant higher-confidence clinical-evidence tier. Relative-frequency estimates were occasionally large because the pooled remainder contained only one alternate allele; therefore, absolute counts, callable allele denominators and exact uncertainty intervals are essential (Table 2). The FTCD chr21:46,154,176 finding was borderline after multiple-testing correction (q=0.0497 across 694 tests), and no observed contrast should be interpreted as an ancestry-specific disease-risk association.

**Table 2.** Six exploratory AFR-elevated pathogenic/likely pathogenic chromosome 21 SNVs.

| Gene / position | Change | Stars | AFR AC / AN | Rest AC / AN | AFR/R | BH q |
|---|---|---|---|---|---|---|
| FTCD 46,137,006 | A>T | 2 | 123 / 33,478 | 19 / 1,427,652 | 276.1 | 1.28×10<sup>−176</sup> |
| DONSON 33,586,090 | A>G | 2 | 10 / 33,480 | 1 / 1,428,382 | 426.6 | 7.41×10<sup>−14</sup> |
| TSPEAR 44,529,799 | A>C | 2 | 8 / 33,474 | 1 / 1,427,910 | 341.3 | 9.27×10<sup>−11</sup> |
| FTCD 46,154,176 | G>A | 2 | 9 / 33,476 | 111 / 1,427,022 | 3.46 | 0.0497 |
| PCNT 46,355,576 | C>A | 1 | 7 / 33,480 | 1 / 1,428,344 | 298.6 | 3.00×10<sup>−9</sup> |
| PCNT 46,443,808 | A>T | 1 | 6 / 33,422 | 1 / 1,425,730 | 256.0 | 9.82×10<sup>−8</sup> |

Note. All positions 1-based GRCh38 chr21; ratio = AFR AC/AN divided by pooled-remainder AC/AN; q adjusted across 694 tests. Group membership is genetically inferred. The pooled remainder is not European or East Asian ancestry alone.

The AFR and pooled-remainder frequency contrasts are visualized in Fig 2.

**Fig 2.**
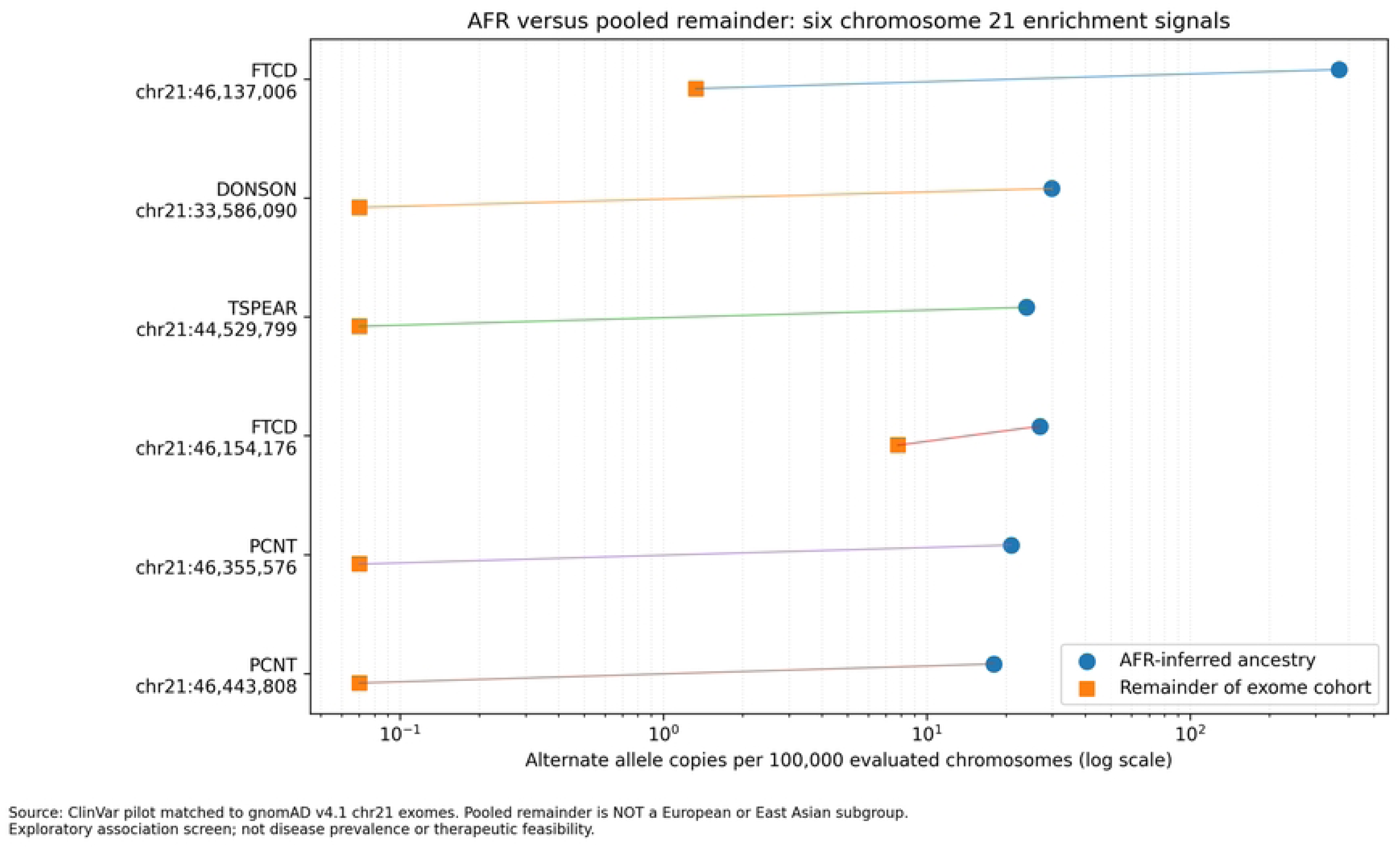
AFR and pooled-remainder frequencies at six exploratory lead variants. Observed alternate-allele frequencies for six exploratory AFR-elevated variants in the AFR subset and pooled remainder of gnomAD v4.1 exomes. Population labels refer to the actual matched exome groups; comparisons should not be interpreted as disease-prevalence estimates.

Under a minimum AFR AC threshold of 3, 5 and 10 (with q<0.05 and ratio ≥2), 13, 6 and 2 variants, respectively, met the rule in the full 694-test family. Thus the selected “elevated” set was sensitive to the allele-count threshold. Exact binomial confidence intervals and the full 694-site table were retained in the supporting analysis package. Median callable allele numbers across audited sites were 33,472 in AFR and 1,427,360 in the pooled remainder, indicating substantial differences in denominator size. No alternate homozygote was reported among the six leads; this does not establish absence of homozygotes in the underlying populations.

### Sequence accessibility substantially narrows chemistry-only eligibility

All 234 reference alleles were consistent with the chromosome 21 FASTA. NGG/PAM-window screening found 118/234 (50.4%) with at least one nominal placement: 105/194 (54.1%) ABE-class and 13/40 (32.5%) CBE-class variants. Across editor models there were 261 candidate placements (74 ABE7.10, 169 ABE8e and 18 BE4). The broader ABE8e window generated more nominal placements by design and should not be interpreted as greater in-vivo editing efficiency. Of all 234 variants, 55 (23.5%) had ≥1 nominal zero-bystander option. For the 47 AFR-observed chemistry-compatible variants, 21 (44.7%) had a placement and 10 (21.3%) a zero-bystander option.

Two higher-confidence AFR-elevated variants illustrate the constraints. DONSON chr21:33,586,090 A>G was chemically CBE-compatible, but lacked a modelled NGG/BE4 placement. FTCD chr21:46,154,176 G>A had two ABE8e options, both containing additional adenines inside the nominal activity window (two and three bystanders, respectively). Thus, neither was a zero-bystander lead under the screened conventional editor assumptions. Failure of this particular screen does not imply that other editors, PAM-relaxed variants or prime editing are infeasible.

### Potential bystander consequences and theoretical multi-base products

Among 261 editor-specific placements, 88 had zero potential window bystanders, 82 had one, 64 had two and 27 had at least three. In total, 295 placement-level bystander instances represented 195 distinct genomic substitution pairs. MANE v1.5 annotation yielded 208 transcript-level records. The unique-allele primary classification was 115 missense, 47 synonymous, 17 intronic splice-region, 10 essential splice-acceptor dinucleotide, two candidate start-codon changes, two 5′-UTR and two other intronic changes. Exonic splice-boundary proximity was separately flagged in 19 cases, overlapping other classes; these annotations are location/consequence predictions rather than functional assays. Primary transcript consequence classes appear in Table 3.

**Table 3.**
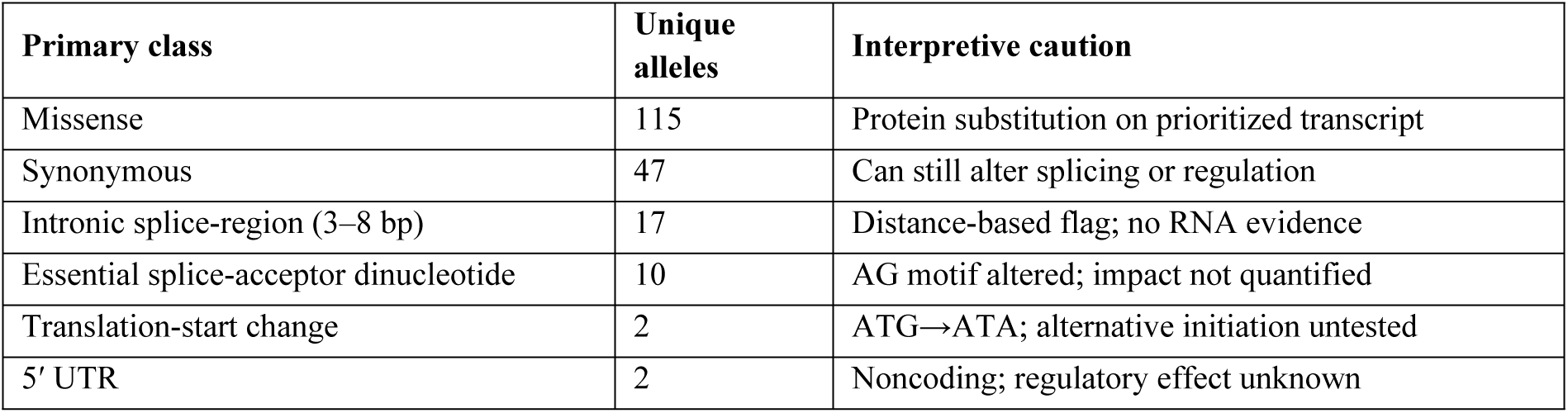
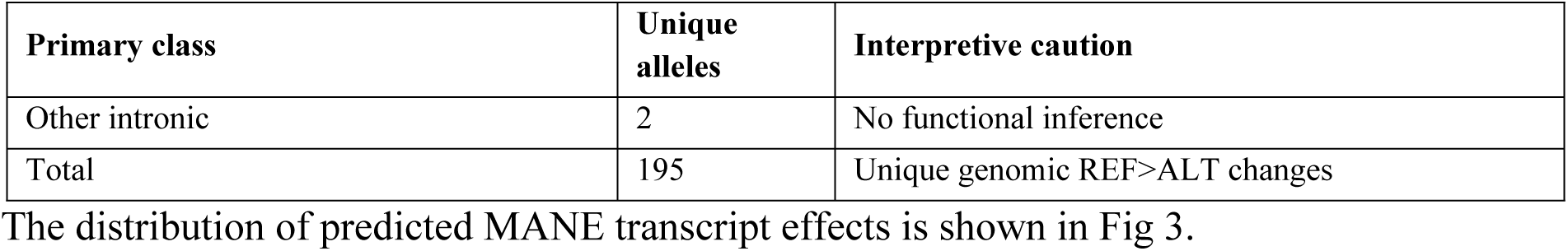
Predicted MANE primary consequences among 195 distinct hypothetical bystander alleles.

| Primary class | Unique alleles | Interpretive caution |
|---|---|---|
| Missense | 115 | Protein substitution on prioritized transcript |
| Synonymous | 47 | Can still alter splicing or regulation |
| Intronic splice-region (3–8 bp) | 17 | Distance-based flag; no RNA evidence |
| Essential splice-acceptor dinucleotide | 10 | AG motif altered; impact not quantified |
| Translation-start change | 2 | ATG→ATA; alternative initiation untested |
| 5′ UTR | 2 | Noncoding; regulatory effect unknown |
| **Primary class** | **Unique alleles** | **Interpretive caution** |
| Other intronic | 2 | No functional inference |
| Total | 195 | Unique genomic REF>ALT changes |

The distribution of predicted MANE transcript effects is shown in Fig 3.

**Fig 3.**
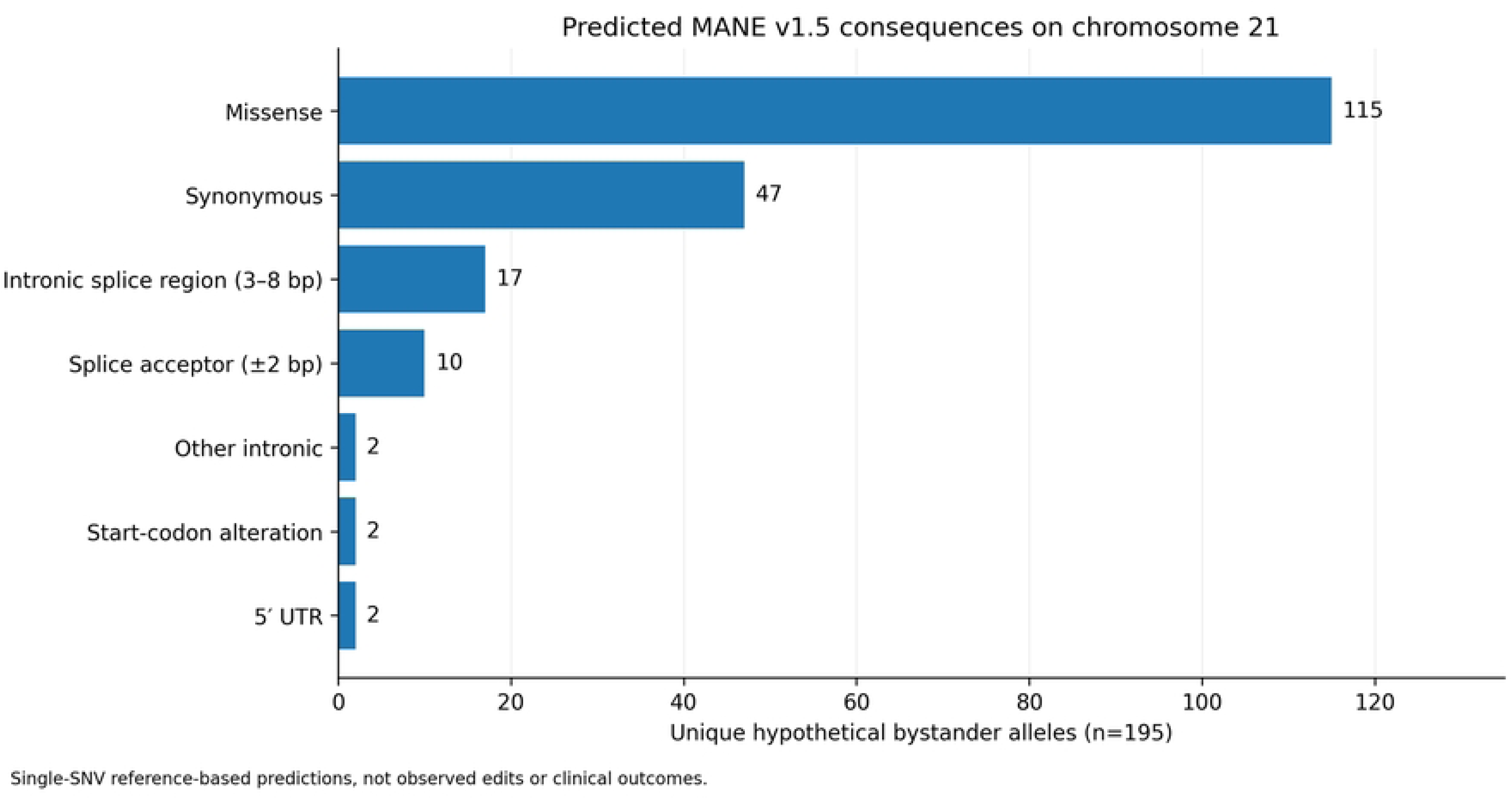
Predicted primary MANE consequences of hypothetical bystander alleles. Primary MANE v1.5 consequence classes assigned to 195 distinct hypothetical bystander substitutions. These are in-silico effects of possible edits, not observed editing products or measured probabilities.

Five unique hypothetical bystander alleles exactly overlapped pathogenic/likely pathogenic ClinVar alleles in the study snapshot: APP p.Ile716Val; RUNX1 p.Asp198Asn and p.Arg162Gly; TMPRSS3 p.Glu104Gly; and AIRE p.Met1Ile at an annotated initiation codon. The two RUNX1 alleles carried expert-panel evidence, motivating follow-up review but not proving harm from editing. In a separate motif audit, all ten predicted essential acceptor variants altered a canonical AG and both translation-start variants altered annotated ATG to ATA. The union of these structural or ClinVar flags comprised 16 distinct alleles associated with 24 guide placements.

Exhaustive binary enumeration across 261 placements produced 1,528 theoretical states, with 764 conditional on the intended target correction. Of these 764, 261 contained no additional bystander and 503 contained ≥1. There were 361 corrected states with at least one modelled MANE amino-acid change, 37 with a prioritized structural/ClinVar flag, and 17 with an exact ClinVar P/LP overlap; the categories overlap. Twenty-six placements contained multiple editable bases in the same codon and 56 corrected states combined such edits. For FTCD p.Arg71Ter, two ABE8e placements yielded 12 conditional corrected combinations, illustrating how a synonymous single change and a missense single change may combine into a different codon-level outcome. No probability model was fitted, so the percentages of enumerated states are not expected product frequencies.

### Guide homology and final candidate filtering

The chromosome 21–restricted homology census examined 187 unique 20-nt guide sequences against 4,106,000 NGG-associated protospacer sites on both strands of the 46,709,983-base reference. Excluding each intended site, 148/187 (79.1%) guides had no additional chromosome 21 NGG site with ≤3 mismatches and 39/187 (20.9%) had ≥1. Across distinct guide–site pairs, 69 additional matches were found (nine with one mismatch, seven with two and 53 with three). The reference-limited, same-placement zero-bystander and no-other-chr21-site criterion retained 70 editor-specific placements corresponding to 44 unique variants, including nine observed in AFR. No whole-genome off-target risk estimate was generated. The chromosome 21 guide near-match distribution is shown in Fig 4.

**Fig 4.**
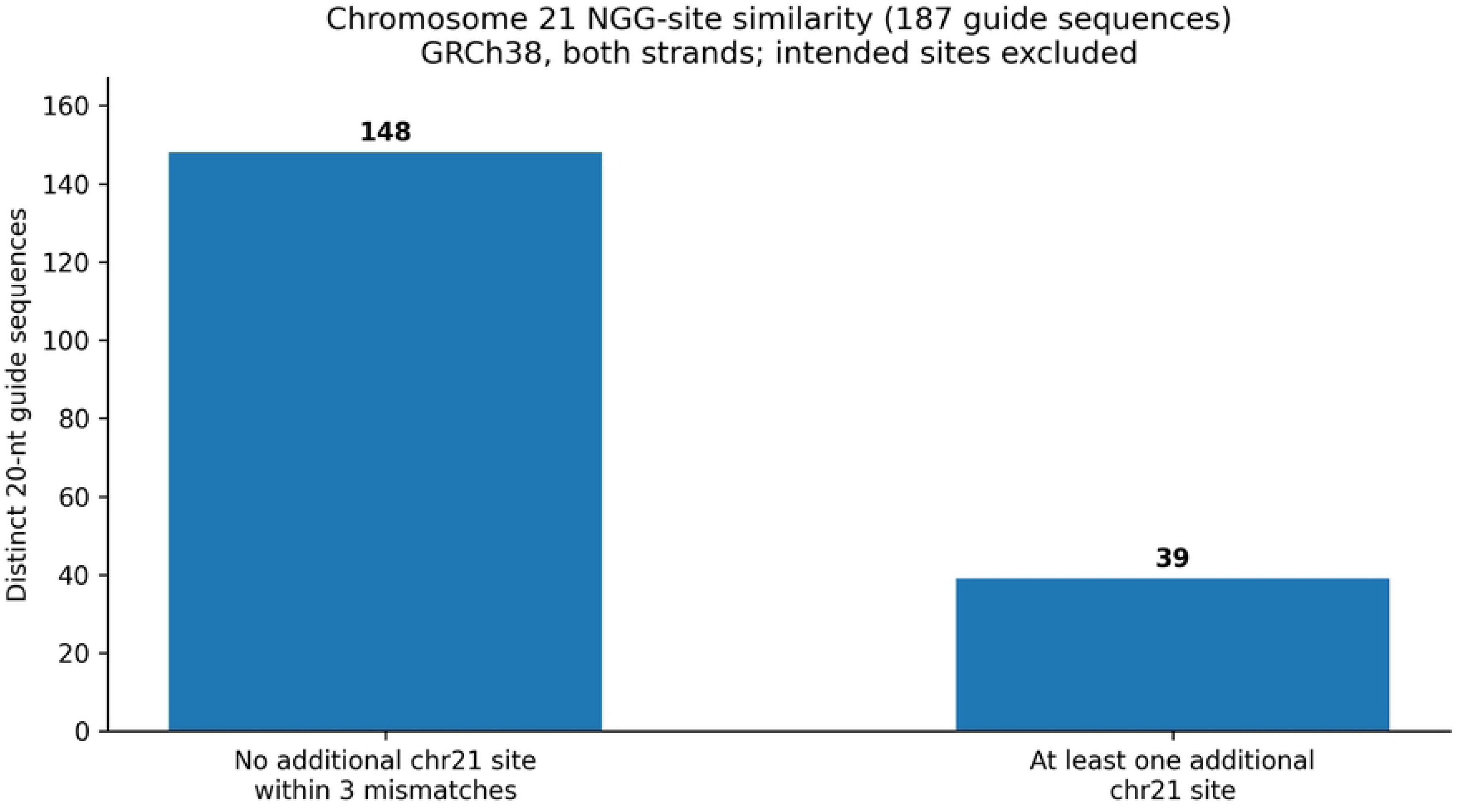
Chromosome 21 near-match sites for 187 candidate guide sequences. Chromosome 21–restricted NGG guide-homology classification of 187 distinct protospacers allowing 0–3 mismatches after intended-site exclusion. Absence of a within-chromosome site is not evidence of genome-wide off-target safety.

### Integrated AFR-observed versus AFR-unobserved comparison

Among the 234 chemistry-compatible primary variants, 47 had AFR AC>0 and 187 had AFR AC=0. No statistically significant contrast was observed for any PAM/window access, any zero-bystander placement or any same-placement chr21-only low-homology plus zero-bystander option (Table 4; all BH q=1.0). Estimates were imprecise, with confidence intervals compatible with meaningful differences in either direction. The observed absence of a statistically detectable difference is therefore not an equivalence result. Of the nine AFR-observed variants meeting the final local low-flag criterion, five had AFR AC=1; all nine were ABE-class. None of the six exploratory AFR-elevated leads passed the final combined local criterion.

**Table 4.**
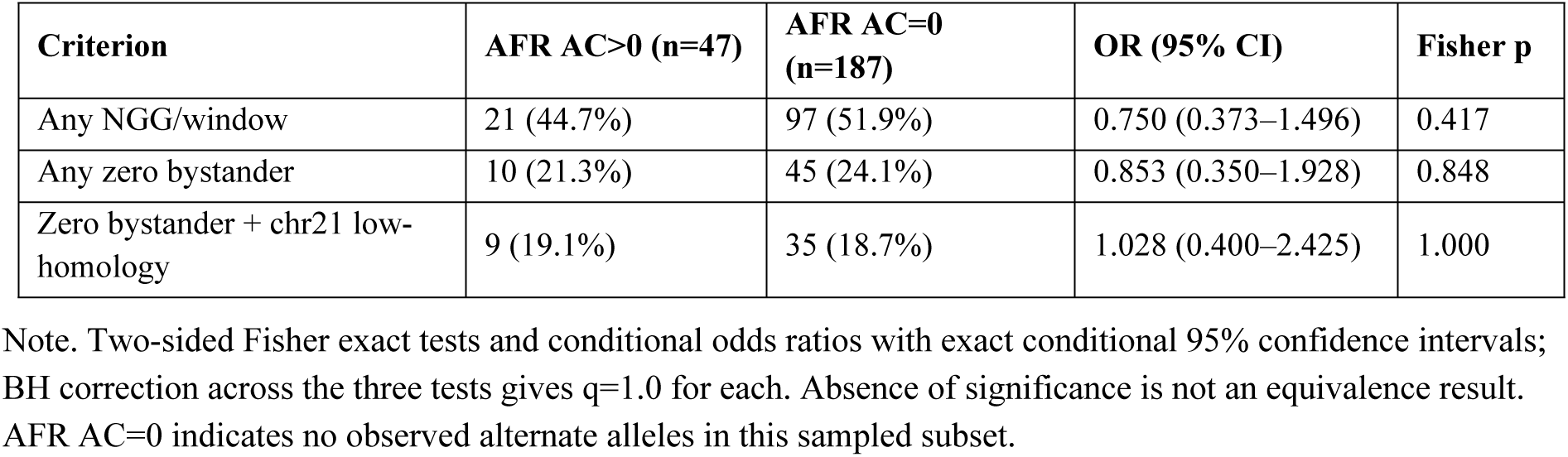
Variant-level editing-accessibility outcomes by AFR observation.

The three accessibility comparisons are shown with exact confidence intervals in Fig 5.

**Fig 5.**
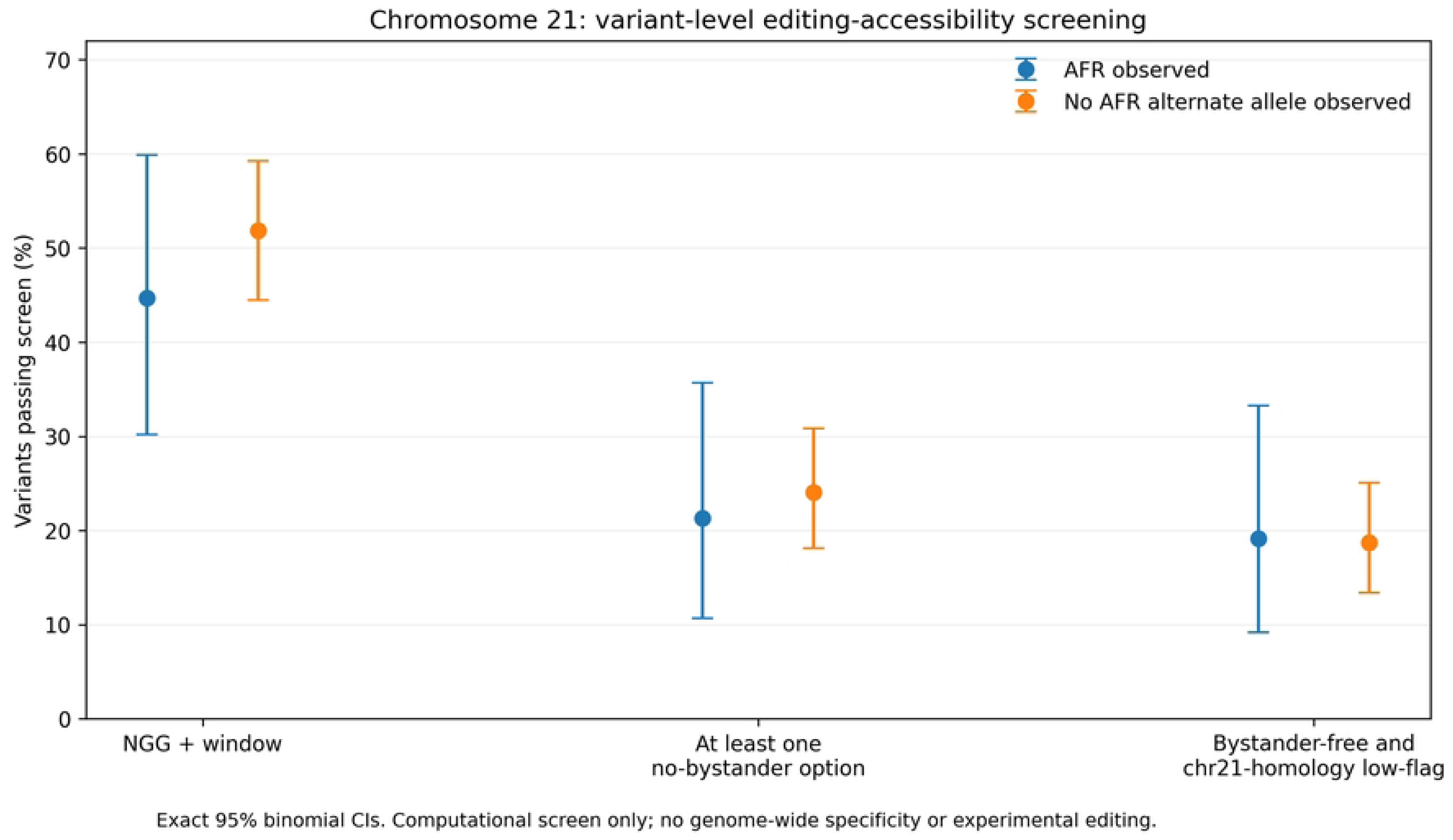
Base-editing accessibility by observed AFR allele status. Three variant-level sequence-accessibility proportions by AFR observation status, with exact binomial confidence intervals. “AFR AC=0” denotes no observed alternate allele in the specified gnomAD AFR exomes, not absence from African populations.

### Allele-count precision and ascertainment-dependent enrichment

The population-frequency screen retained 694 overall-gnomAD-observed, PASS variants and found 109 with at least one alternate allele in the gnomAD AFR subset. Median called-allele number was 33,472 for AFR and 1,427,360 for the pooled remainder, demonstrating an approximately 43-fold disparity in available denominators. The large imbalance means that very high AFR/remainder ratios may coincide with wide uncertainty in the comparator estimate, particularly when the remainder contains only one alternate allele. Across the 694 audited variant records, 21 had a reported alternate-allele homozygote in the overall cohort, whereas none had a reported alternate-allele homozygote in the AFR subset. None of the six flagged AFR-elevated variants had an alternate-allele homozygote in either cohort.

At the prespecified minimum of five AFR alternate alleles, six variants fulfilled the exploratory elevation rule. Relaxing the minimum to three increased the number of flags to 13, while requiring at least 10 reduced it to two. The retained strongest signals under the most stringent count rule were FTCD chr21:46,137,006 and DONSON chr21:33,586,090. These thresholds are analytically transparent but cannot correct ascertainment bias in ClinVar submissions or distinguish the heterogeneous populations represented by the gnomAD grouping. Notably, alleles with zero AFR counts were “not observed in the sampled subset” rather than “absent from Africans” [7,24–27]. Additional ascertainment and threshold findings are summarized in Table 5.

**Table 5.** Additional population-uncertainty and threshold-sensitivity findings.

| **Metric** | **Observed result** | **Interpretation** |
|---|---|---|
| AFR-observed among 694 PASS sites | 109/694 (15.7%) | Occurrence within sampled AFR exomes |
| Median AFR allele number | 33,472 | Different denominators across ancestry groups |
| Median pooled-remainder allele number | 1,427,360 | Not a specific EUR/EAS comparator |
| Variant records with Alt/Alt overall | 21/694 | Not frequency of affected individuals |
| Variant records with Alt/Alt AFR | 0/694 | Not proof of population absence |
| Elevated flags, AFR AC ≥3 | 13 | Exploratory cutoff sensitivity |
| Elevated flags, AFR AC ≥5 | 6 | Prespecified screen |
| Elevated flags, AFR AC ≥10 | 2 | Most stringent count screen |

Note. All figures refer to the restricted study dataset; confidence bounds for individual allele frequencies are in the accompanying Stage 8 records.

The precision of the six lead-variant frequency estimates is visualized in Fig 6.

**Fig 6.**
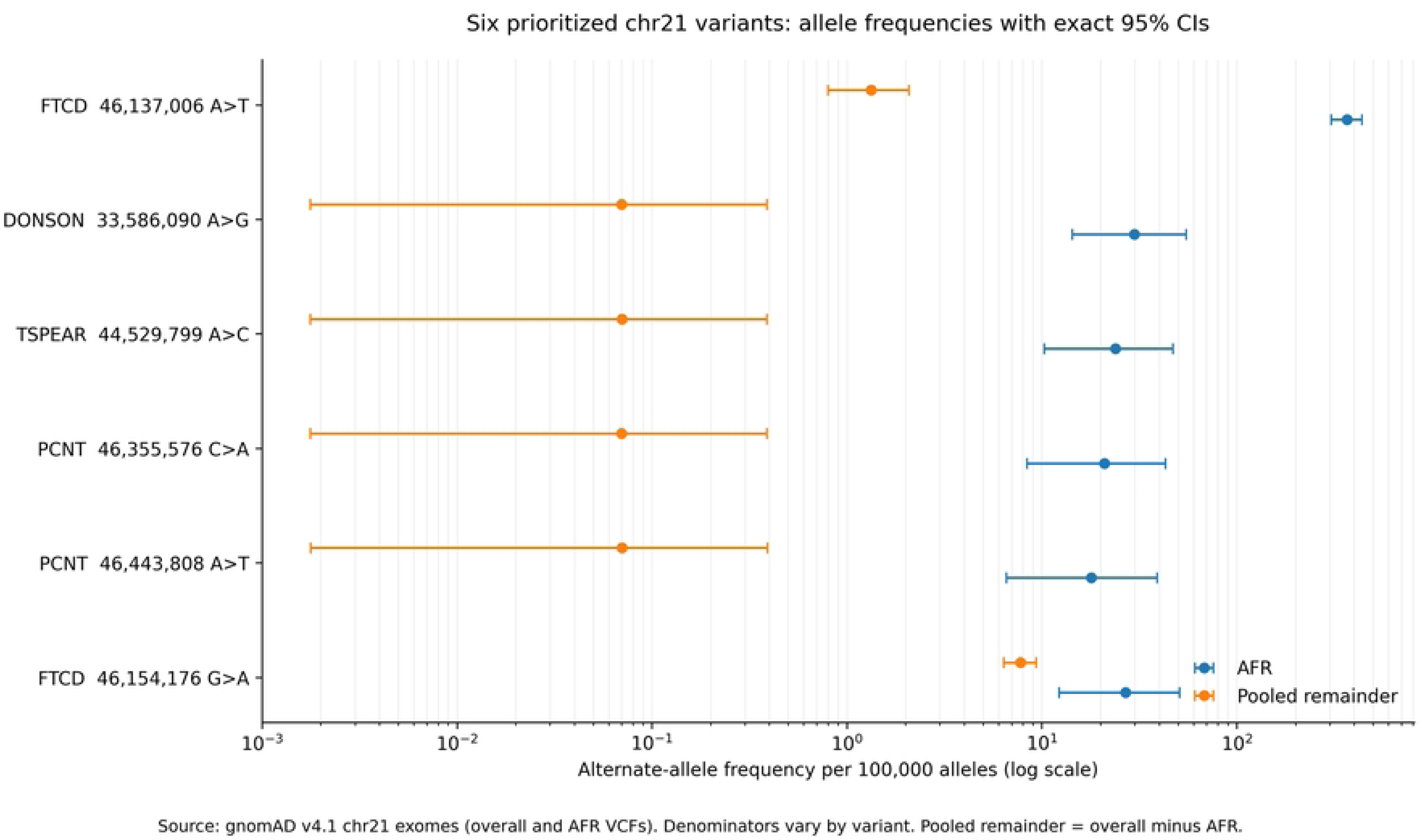
Exact confidence intervals for population allele-frequency estimates. Exact 95% binomial confidence intervals for observed AFR and pooled-remainder alternate-allele frequencies at six exploratory AFR-elevated chromosome 21 variants. The figure uses gnomAD allele counts and is not an estimate of disease penetrance or prevalence.

### Quantitative distribution of potential bystanders and clinical annotation flags

Across 261 editor-specific placements, 88 had no nominal editing-window bystander, 82 had one, 64 had two, and 27 had three or more. These placement-specific observations corresponded to 295 potential bystander conversion events but only 195 distinct genomic substitutions, because guide placements and editor models can interrogate overlapping positions. By editor architecture, the 295 event instances were distributed as 29 for ABE7.10, 245 for ABE8e and 21 for BE4; the larger ABE8e count follows directly in part from its broader assumed window and should not be interpreted as higher measured off-target toxicity.

Reference-locked MANE v1.5 annotation assigned 115 of the 195 distinct alleles to a primary missense class, 47 to a synonymous class, 17 to an intronic splice-region proximity class, 10 to essential splice-acceptor dinucleotide changes, two to altered translation-initiation codons, two to 5′-untranslated regions and two to other intronic sequence. The 195 alleles yielded 208 transcript-specific records because some alleles intersect more than one MANE model. Nineteen additional splice-boundary proximity annotations were recorded as overlapping flags, not additional unique variants.

The clinical cross-reference found five distinct hypothetical substitutions that matched ClinVar pathogenic/likely pathogenic alleles, occurring in eight placement instances. Two are expert-panel-reviewed RUNX1 alleles; the others map to APP, TMPRSS3 and AIRE. Their match is a clinically meaningful flag for follow-up, but the database classification pertains to a variant– phenotype interpretation and does not prove that an editor generates that product or that its effect in a co-edited haplotype is equivalent to its isolated clinical allele [18,20]. The matching ClinVar alleles are listed in Table 6.

**Table 6.** Potential ClinVar-overlapping bystander products on chromosome 21.

| **Gene** | **Position (GRCh38)** | **Hypothetical change** | **MANE consequence** | **ClinVar review** |
|---|---|---|---|---|
| APP | 25,891,787 | T>C | p.Ile716Val | 1 star |
| RUNX1 | 34,859,495 | C>T | p.Asp198Asn | 3 stars |
| RUNX1 | 34,880,581 | T>C | p.Arg162Gly | 3 stars |
| TMPRSS3 | 42,388,940 | T>C | p.Glu104Gly | 1 star |
| AIRE | 44,286,009 | G>A | p.Met1Ile; start altered | 1 star |

Note. These exact alleles were cross-referenced against the study ClinVar snapshot, and are theoretical bystander outcomes; clinical effect, editing frequency and allelic phase were not tested.

The ClinVar overlap flag counts are visualized in Fig 7.

**Fig 7.**
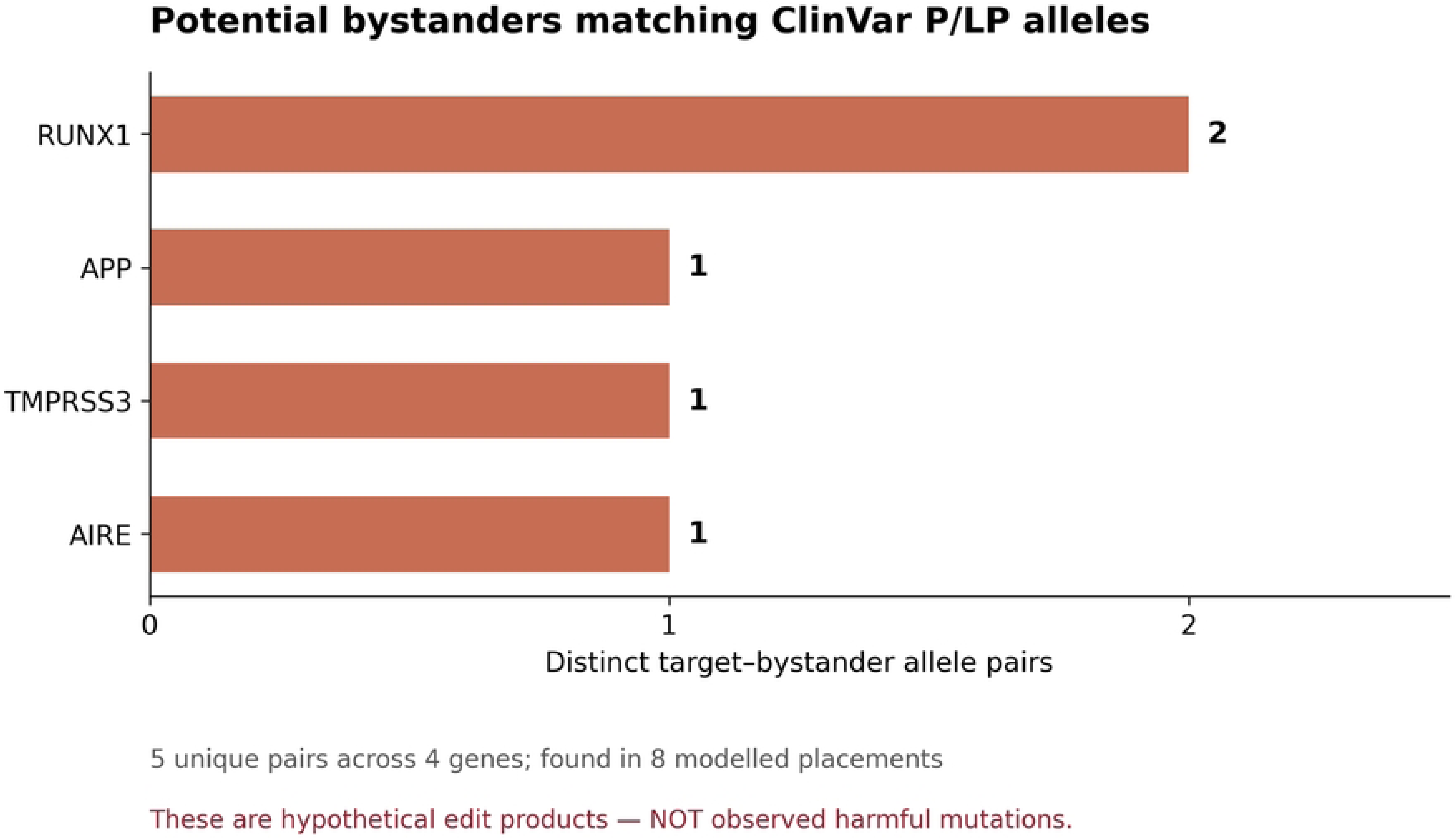
ClinVar overlap flags among theoretical bystander substitutions. Distribution of bystander substitutions with direct pathogenic/likely pathogenic ClinVar allele overlap compared with other theoretical bystander products. The analysis is a clinical annotation flag, not an observed editing event.

A separate structural audit produced a nonredundant set of 16 distinct bystander alleles meriting manual review: ten substitutions change canonical splice-acceptor AG motifs, two alter annotated ATG initiation codons, and five exactly overlap ClinVar pathogenic/likely pathogenic records, with one AIRE allele satisfying both start-codon and clinical-overlap criteria. The 16 alleles were represented in 24 of the 261 placements; only three were associated with an editing target observed in the AFR subset. The overlap among categories, and the distinction between the target variant and the hypothetical bystander, are essential to any downstream biological interpretation [13,34].

### Enumeration of joint on-target and bystander edited sequence states

The 261 placement-specific bystander sets generated 1,528 enumerated binary sequence states when both target-retained and target-corrected branches were included. Half of these—764 states—were conditional on the intended ALT→REF correction, comprising 261 target-only configurations and 503 additional configurations with one or more bystander substitutions. Among the 764 corrected states, 361 had at least one predicted MANE amino-acid change relative to the reference; 37 included a high-priority structural/clinical bystander allele; 17 contained an exact ClinVar P/LP allele; 19 included a canonical splice-acceptor candidate; and three included a start-codon change. These categories overlap. Their enumeration does not imply that such molecular products occur with equal probability, or at all, in a particular cell type.

The dependence between nucleotide edits becomes especially clear when distinct editable bases occupy the same coding triplet. Twenty-six placements contained at least two bystander sites in a shared MANE codon and 56 theoretical target-corrected configurations contained multi-bystander same-codon changes. Because the amino acid is translated from the joint codon, the consequence of two substitutions must not be inferred as the sum of independent single-substitution annotations. Quantitative product-mixture modelling would require experimental measurements or validated sequence-conditioned outcome models [11,12,35]. The analytical units and configurations are defined in Table 7.

**Table 7.**
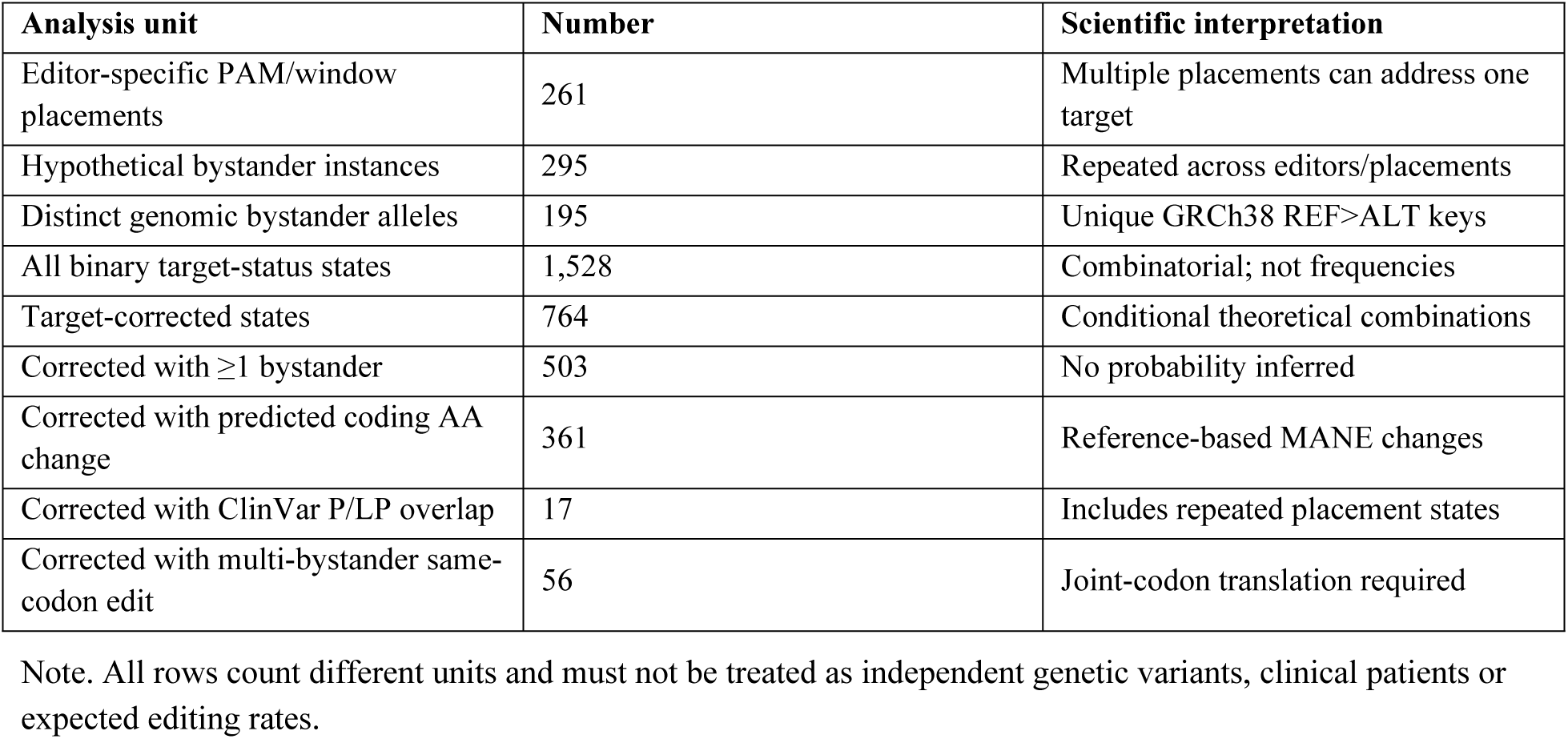
Distinct analysis units and combinatorial outcomes.

The combinatorial editing-state counts are illustrated in Fig 8.

**Fig 8.**
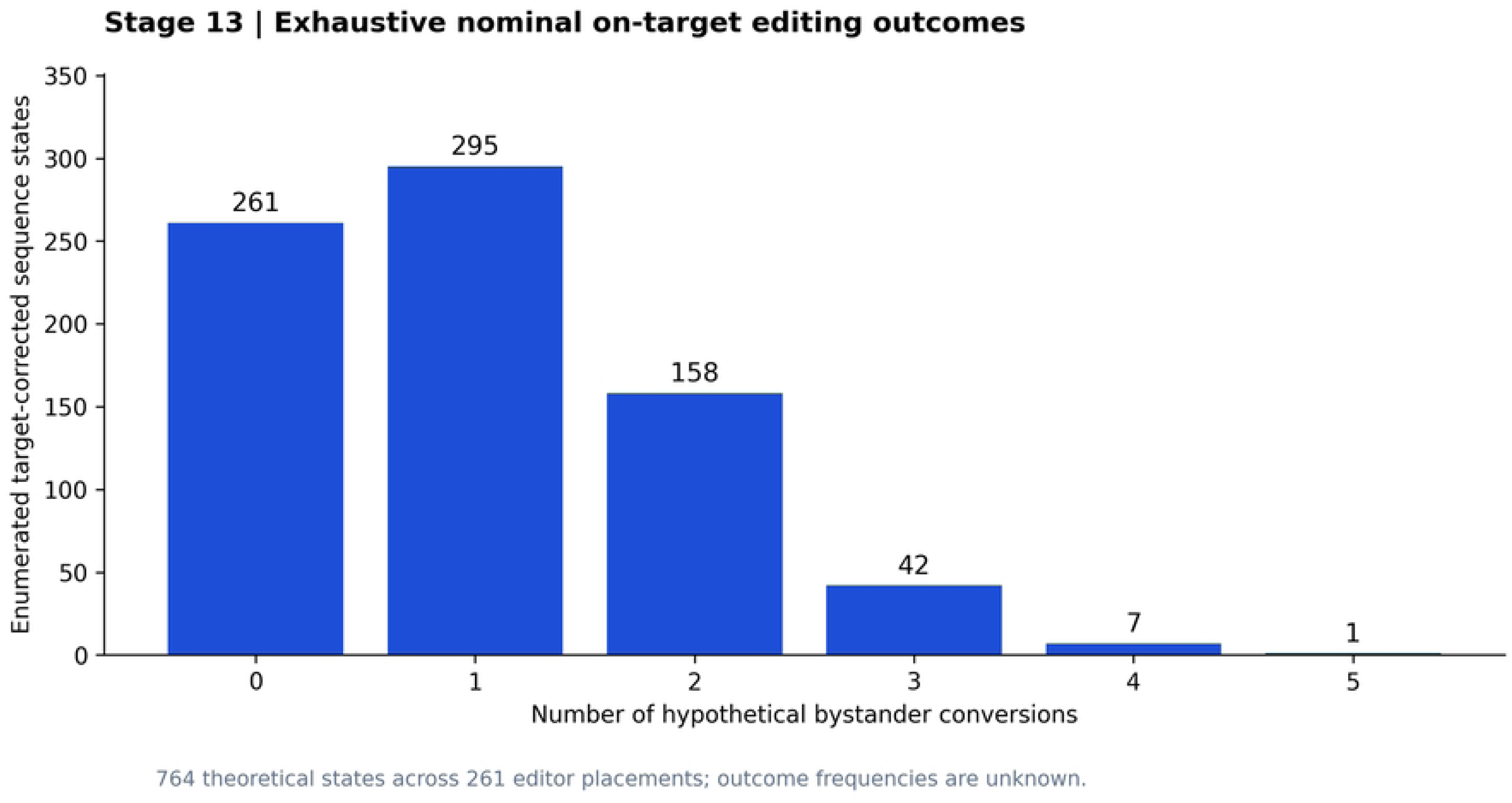
Numbers of theoretical outcomes with accompanying bystander edits. Theoretical target-corrected editing-outcome configurations classified by numbers of accompanying bystander edits. Counts enumerate logical subsets under the nominal window model and do not estimate edit-product abundance.

### Target-specific illustration: DONSON and FTCD

DONSON chr21:33,586,090 A>G (ClinVar two-star; AFR AC=10, AN=33,480) illustrates why ancestry-related interest alone does not ensure canonical base-editor accessibility. Although the implied ALT→REF correction has CBE-compatible chemistry, no suitable BE4/NGG PAM-window placement was found in the specified model. This negative finding is editor-architecture-specific: it does not exclude altered-PAM systems, different window configurations, prime editing or other modalities. It should motivate comparative tool selection rather than a categorical assertion that the allele is uneditable [23,38,41].

FTCD chr21:46,154,176 G>A (two-star ClinVar; AFR AC=9/33,476; AFR/remainder frequency ratio=3.46) yielded two nominal ABE8e/NGG placements. Both lacked additional chromosome 21 NGG sites within the restricted 0–3 mismatch homology search, yet both had potential window bystanders (two and three, respectively). The two placements together produced 12 conditional target-corrected bystander-subset configurations. Their transcript-aware interpretation identified the possible p.Leu72Pro and p.Ile73Thr consequences, depending on which nearby adenines are altered. At chr21:46,154,171 A>G individually, the putative effect is p.Leu72Leu; chr21:46,154,172 A>G individually produces p.Leu72Pro; both substitutions together also translate as p.Leu72Pro. This case demonstrates the importance of modelling molecular products and target-rescue consequences jointly, rather than declaring a target successful because its pathogenic nucleotide can be reverted.

The second FTCD signal, chr21:46,137,006 A>T, is not compatible with the canonical single-step ABE/CBE correction chemistry screened here. On the MANE Select transcript it is annotated as a near-terminal stop variant, p.Leu536Ter, whereas the chemically compatible p.Arg71Ter variant represents an early premature termination event. These differing transcript contexts mean that the two FTCD alleles should not be conflated in functional interpretation, even though both were among the exploratory AFR-elevated list. Neither the reference sequence nor the population data establish the fraction of function that must be restored for clinical benefit.

### Integrated local low-flag shortlist and evidentiary caveats

The final chromosome 21-limited gate selected 70 guide/editor placements corresponding to 44 distinct higher-confidence, chemistry-compatible target variants with no additional deaminatable base inside the modelled editing window and no other chromosome 21 NGG protospacer with ≤3 substitutions. Only nine of these 44 target variants were observed in the AFR exome subset, and all nine were in the ABE chemistry class. Five of the nine had an AFR alternate-allele count of one. Consequently, the shortlist is best described as locally uncomplicated under the chosen editor models, not as a set of strongly AFR-enriched, safe or clinically prioritized treatments. The nine AFR-observed variants are listed in Table 8.

**Table 8.**
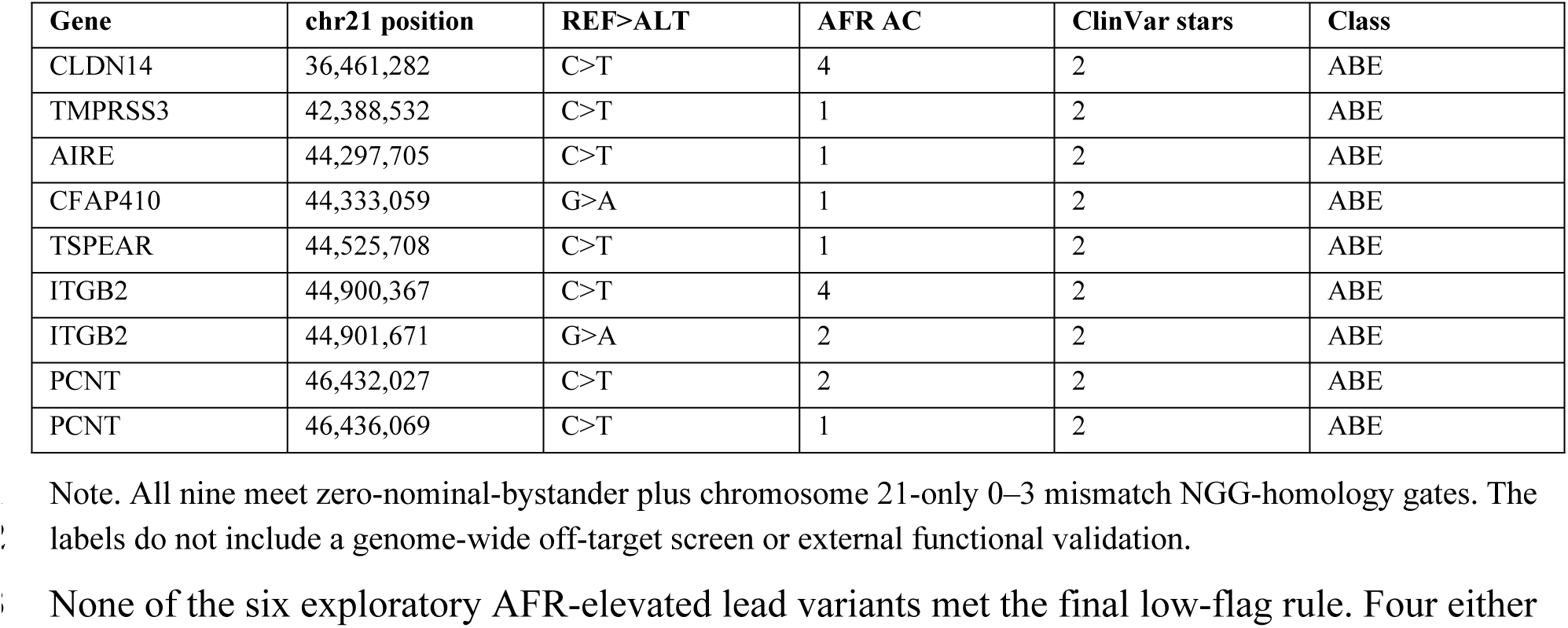
Nine AFR-observed target variants satisfying the limited local low-flag screen.

None of the six exploratory AFR-elevated lead variants met the final low-flag rule. Four either lacked canonical single-step base-editing chemistry or failed the high-confidence chemistry screening definition; DONSON had no modelled NGG/BE4 placement and FTCD p.Arg71Ter had bystander bases at every modelled ABE8e placement. Thus, the two clinical-priority concepts—population frequency elevation and a locally simple editing design—identified substantially different subsets. This divergence is a central finding rather than an analysis failure: molecular implementability cannot be inferred from enrichment alone.

### Internal validation and analysis boundaries

Source-reference integrity checks confirmed the REF nucleotide for all 234 chemistry-compatible candidate variants against GRCh38 chromosome 21 and for all 295 hypothetical bystander-placement events. The downloaded MANE GRCh38 v1.5 annotation was confirmed using its published checksum; all 215 chromosome 21 MANE transcript models used in the relevant analysis passed coding-sequence phase and annotated initiation-codon consistency checks. An independent in-house consequence implementation produced no discrepancies in the tested alleles. A streaming refactor of the chromosome 21 guide scanner reproduced all 256 0–3 mismatch guide–site tuples, including 187 intended reference loci and 69 additional chromosome 21 near-matches. Neither an independent release-matched Ensembl VEP execution nor a full primary-genome specificity scan was completed; such external validation is still required before interpretation of these candidates as therapeutically acceptable.

## Discussion

This pilot demonstrates the value of decomposing the phrase “genome-editing amenability” into successive, auditable questions. A chemical ALT→REF transition was possible for 57.1% of the original reviewed ClinVar chromosome 21 SNVs. Yet, after requiring higher-confidence clinical status, observed overall gnomAD data, a conventional NGG PAM, a nominal editor window and bystander-focused filters, a substantially smaller set remained. The step from chemical class to 118 nominally targetable variants—then to 55 nominally zero-bystander variants—captures constraints that would be invisible in a chemistry-only survey. Importantly, these are *model-defined* attrition steps; narrower windows or alternative editors could yield different eligible sets [1–4,10–12].

The ancestry component is hypothesis-generating rather than definitive. Six variants met the AFR-elevation screening rule, but their selection depended on a minimum alternate-allele count, and large frequency ratios were sometimes driven by a single comparison-cohort allele. The gnomAD AFR group aggregates heterogeneous ancestries, and the pooled remainder includes multiple other genetic-ancestry groups with different sample contributions. Our results cannot support a specific European-versus-East-Asian contrast, claims about African national populations, or clinical disease prevalence. Stronger biological conclusions require independent cohort replication, site-level ascertainment review, phenotype and segregation evidence, and finer-scale genetic-ancestry analyses [6–9,19].

The lack of a statistically significant association between AFR observation and three nominal accessibility outcomes is informative in a restricted sense: this analysis does not support assuming that variants observed in the sampled AFR subset are intrinsically more or less compatible with the screened editors. The wide odds-ratio confidence intervals limit the conclusions, and the two groups were selected conditional on ClinVar evidence, overall exome observation and chemistry. Such selection could induce differences unrelated to ancestry, while regional sequence context and variation in clinical ascertainment remain potential confounders. The analysis was not designed or powered as an equivalence study.

The bystander and theoretical-outcome results argue against using a single sequence-level “clean edit” label without transcript-aware interpretation. An ostensibly silent bystander can be close to a splice boundary, a motif-altering change can be functionally buffered, and two changes in the same codon may be nonadditive. The five exact ClinVar-overlapping substitutions, especially expert-panel RUNX1 alleles, provide concrete examples of why potential editing products should receive independent review. They do not demonstrate that a given editor generates those alleles or that a specific edit would cause a disease phenotype. Joint-product modeling and sequencing of actual editing outcomes would be needed to establish such effects [11–13,18].

The distinction between local homology and whole-genome specificity is equally important. Although 148 of 187 unique guides lacked an additional chromosome 21 NGG-associated site within three mismatches, that finding says little about the other chromosomes, noncanonical PAMs, DNA bulges, cell-specific chromatin or deaminase-associated off-target activity. The resulting 44-variant low-flag set is a reproducible list for follow-up screening, not a list of safety-qualified therapeutic guides. Even the AFR-elevated FTCD p.Arg71Ter lead, which had no additional short-range chromosome 21 matches in this screen, retained nominal bystander opportunities. This illustrates how specificity dimensions can conflict [10–12,14].

The pilot has practical value as a transparent data-processing template. Each analysis retains raw source identity, matching coordinates, threshold definitions, denominator sizes and level of evidence; coding and transcript consequences are tied to a named reference build and versioned MANE annotation. Expansion to the autosomes should reuse the same evaluation definitions but not simply pool results without chromosome-wise coverage and ancestry-structure controls. A clinically informed prioritization model would further integrate inheritance, allelic dosage, age of onset, tissue accessibility, target-cell delivery and disease-mechanism-specific functional rescue.

### From chemical permissiveness to biological plausibility

The central conceptual contribution of this study is that genome-editing feasibility cannot be summarized by a single denominator or threshold. More than half of the reviewed ClinVar alleles were chemically reversible through one of two standard base-editor reaction classes, yet among 234 higher-confidence, observed chemistry-compatible alleles, only half met an NGG/window rule and fewer than one-fifth retained a locally uncomplicated same-placement option. Each narrowing step addresses a different biological question, and each imposes assumptions about the editor, allele sequence and intended product. The output is therefore a hierarchy of evidence, not an ordered probability of treatment success [1–4,22,23].

At the gene-therapy level, rescuing the relevant phenotype depends on the gene’s mechanism of disease and the target tissue. Haploinsufficiency, dominant-negative and gain-of-function disorders demand distinct approaches; even a technically clean ALT→REF correction may be insufficient in a tissue with inadequate delivery, irreversibly damaged cells or a requirement for near-complete correction. Conversely, a partial correction fraction may be highly consequential for a disease with a selective growth advantage in corrected cells. The successful clinical development of hematopoietic gene-editing therapies and early examples of patient-specific in-vivo editing should be understood in that tissue- and disease-specific framework, rather than generalized to every coding variant [28–30,39,40].

The negative DONSON finding illustrates an important methodological asymmetry: absence of a canonical NGG/BE4 design under our particular geometry cannot justify rejecting the underlying disease allele as an editing target. Engineering alternative PAM recognition, changing editor window geometry or employing prime editing may overcome the observed constraint, although each alternative demands its own assessment of product purity, efficiency, delivery and specificity. It is scientifically preferable to label this candidate “not accessible under the tested model” than “not editable” [4,23,37,38,41].

### Clinical heterogeneity concealed by allele-level labels

ClinVar record-level pathogenicity offers a valuable starting filter but is not equivalent to a uniformly strong, condition-specific causal claim. Our separation of 2–4-star and one-star records demonstrates that evidence tiers vary materially in their composition of nucleotide chemistry. In addition, classifications can reflect different phenotypic contexts or modes of inheritance; careful investigation should therefore return to the relevant RCV/SCV evidence and expert disease-gene validity, particularly for the secondary one-star PCNT variants [5,18,20].

The two FTCD premature-stop alleles are a useful illustration. An early p.Arg71Ter alteration and a near-terminal p.Leu536Ter alteration cannot automatically be assigned the same severity or functional mechanism; nonsense-mediated decay depends in part on transcript architecture, and the residual function of a truncated protein cannot be settled by a stop-gain label alone. Similarly, a hypothetical AIRE initiation-codon bystander is a reason to assess alternative start sites and protein expression, not proof of absent AIRE translation. Precision therapeutics must ultimately connect the predicted genomic product to gene expression, biochemical function and disease phenotype.

### Ancestry representation, sampling uncertainty and equitable translation

The six exploratory AFR-elevated sites identify variants deserving further investigation in well-characterized cohorts; they do not establish population-specific disease susceptibility or identify a pan-African therapeutic portfolio. African genetic ancestry is deeply structured, and the aggregate AFR category of a global exome resource cannot substitute for geographically and ethnolinguistically granular sequencing, appropriate control ascertainment or clinical phenotype linkage [7–9,24–27]. The very unequal called-allele denominators here, and the dependence of the shortlisted signals on minimum alternate-allele thresholds, reinforce the need for population-aware but statistically conservative interpretation.

The null variant-level comparison of AFR-observed and AFR-unobserved accessibility should be interpreted equally carefully. Across the three nominal outcomes, the estimated odds ratios were close to or below one but confidence intervals were broad, particularly for the stricter filters. These are non-significant exploratory associations in a highly selected cohort of ClinVar-submitted, overall-gnomAD-observed alleles; they are not equivalence tests and do not address differences in actual patient-editing efficiency. Nor is it appropriate to infer that the underlying rates of potentially editable disease mutations are identical between ancestry groups [8,24–27].

### Product purity, specificity and experimental triage

A practical guide-development pipeline should prioritize molecular product resolution rather than the binary existence of a PAM. The 261 placement screen identified 295 hypothetical bystander instances, and the 764 corrected sequence configurations illustrate the breadth of plausible alleles even before one accounts for off-window edits, indels and editing kinetics. Protein annotations are particularly important for changes clustered in one triplet, but they must be combined with experimental product frequencies to determine whether an unwanted edit is rare, common or nearly obligatory. High-throughput base-editor studies provide valuable priors for such work, but cannot replace target- and cell-specific testing [11,12,22,35].

Guide homology adds another orthogonal axis: an in-chromosome low-homology result can coexist with an undesirable local bystander profile, as demonstrated for FTCD p.Arg71Ter. Moreover, our chromosome 21 index was intrinsically incapable of finding off-target-like sites elsewhere in the genome and did not address guide–DNA bulges, alternative PAMs, off-target RNA deamination, chromatin effects or repair-associated products. Genome-wide guide mapping, deaminase-specific profiling, and orthogonal sequencing-based assays are necessary before a therapeutic specificity claim is defensible [14,31–33,35,36].

For a future experimental program, the immediate priority is to validate prioritized transcript consequences with a release-locked independent annotation pipeline and splice-disruption scores; to reconstruct representative patient haplotypes for the selected disease alleles; to compare candidate editors in a disease-relevant cellular model; and to use deep amplicon or long-read sequencing for joint edit products together with appropriate genome-wide off-target assays. These steps are proposed future validation, not part of the present study. The analytical strength of the current pilot is to define which claims are presently supported and which depend on the next evidence tier [17,34,35].

### Limitations and future validation

Clinical evidence and ascertainment. ClinVar review status indexes the level of assertion review rather than validating a disease association in every phenotype context. We analyzed pathogenic/likely pathogenic SNVs that were both represented in ClinVar and observable in the overall exome dataset; absent, poorly covered or ClinVar-unrepresented variants were outside the screening denominator rather than demonstrated uneditable. Condition-specific assertions, inheritance and independent gene–disease validity require further reconciliation [5,18].

Population comparisons. gnomAD provides aggregate called-allele counts with unequal site-level denominators, heterogeneous ancestry labels and no individual-level adjustment for relatedness, cohort sampling, batch effects, admixture or fine-scale substructure. The pooled non-AFR remainder is not a European or East Asian population estimate. Allele-count Fisher tests are descriptive exploratory screens and should not be used to infer disease prevalence, penetrance or ancestry-specific causality. Replication in ancestry-resolved cohorts would be needed before population claims.

Editor models and sequence products. NGG PAMs and nominal ABE7.10, ABE8e and BE4 activity windows are deliberately simplified; alternative editors and context-dependent edit probabilities were not assessed. No-window-bystander status excludes only additional canonical deaminatable bases in these windows, not possible indels, RNA editing or other unintended products. The 1,528 enumerated sequence states represent theoretical possibilities, not a probability distribution, and patient-specific phased haplotypes were unavailable.

Functional interpretation. MANE v1.5 coding and splice-motif annotations were confirmed against the local genomic reference and cross-checked in-house, but no release-locked external VEP run, splice-prediction validation or transcript/protein assay was carried out. ClinVar-overlapping hypothetical bystanders are prioritization flags, not observed edited alleles or proof of deleterious effects.

Specificity and translation. Guide matching was exhaustive only for chromosome 21, 20-nt NGG-associated protospacers and up to three ungapped mismatches; it did not evaluate other nuclear chromosomes, alternative contigs, bulges, divergent sites, chromatin context or guide-specific experimental off-target activity. No cellular editing, product sequencing, therapeutic delivery or safety experiments were performed. The results therefore support computational prioritization, not preclinical efficacy or safety claims [11,12,31–36].

## Conclusions

A population-aware, clinically tiered pipeline reduced 234 higher-confidence chemistry-compatible chromosome 21 variants to 118 with a nominal NGG/editor-window placement, 55 with a nominal zero-bystander option and 44 with a same-placement zero-bystander and chromosome 21–restricted low-homology option. Six variants exhibited exploratory elevation in the gnomAD AFR subset, whereas no statistically significant difference in nominal accessibility was detected between AFR-observed and AFR-unobserved variants within the selected analysis cohort. Transcript-aware bystander auditing and joint-state enumeration highlighted why sequence accessibility cannot be equated with clean or therapeutically successful correction. The resulting framework is reproducible and hypothesis-generating; whole-genome and experimental validation remain necessary before clinical interpretation.

## Declarations

Ethics approval and consent to participate. No individuals were recruited or directly contacted, and the analyses were restricted to publicly released aggregate allele counts and clinical annotation datasets. No identifiable participant-level data were accessed and no new participant consent was collected. Any institution-specific determination required by a target journal should be confirmed before submission.

## Data availability

All data necessary to reproduce the reported numerical findings and figures are provided in the Supporting Information (S1–S10 Datasets), together with analysis scripts, metadata and file checksums (S11 File). The analyses used publicly available ClinVar GRCh38 (file date 1 November 2026; https://ftp.ncbi.nlm.nih.gov/pub/clinvar/vcf_GRCh38/), gnomAD v4.1 exome chromosome 21 overall and African/African-American genetic-ancestry allele-count datasets (https://gnomad.broadinstitute.org/), the UCSC GRCh38 chromosome 21 reference sequence (https://hgdownload.soe.ucsc.edu/goldenPath/hg38/chromosomes/chr21.fa.gz), and MANE GRCh38 v1.5 transcript annotations (https://ftp.ncbi.nlm.nih.gov/refseq/MANE/MANE_human/release_1.5/). No individual-level participant data were accessed. The original public-source resources are not redistributed in the Supporting Information. The code and derived tables support inspection and reproduction of the reported results, subject to the software and source-file requirements detailed in S11 File.

## Acknowledgments

No acknowledgments are reported.

## Supporting information

**S1 Dataset.** All 1,944 reviewed ClinVar chromosome 21 pathogenic/likely pathogenic single-nucleotide records with exact-match and overall/AFR population-frequency annotations. Original ClinVar assertions and source release metadata should be verified against source records.

**S2 Dataset.** The 694 PASS and overall-allele-observed ClinVar–gnomAD variants, with recomputed AFR and pooled-remainder allele counts, frequencies, and multiple-testing results.

**S3 Dataset.** The 234 higher-confidence, overall-gnomAD-observed chromosome 21 variants with canonical ABE/CBE correction chemistry and integrated sequence-screening results.

**S4 Dataset.** The 261 nominal NGG PAM/editor-window guide placements, including predicted bystander positions and annotations.

**S5 Dataset.** The 195 distinct hypothetical bystander substitutions and their MANE GRCh38 v1.5 transcript-consequence annotations.

**S6 Dataset.** The 764 enumerated hypothetical target-corrected sequence outcomes across 261 placements; these are not observed editing products or probabilities.

**S7 Dataset.** All 1,528 conditional theoretical sequence configurations (target corrected or unchanged) across editor placements.

**S8 Dataset.** The 187 distinct 20-nucleotide protospacers and chromosome 21–restricted NGG homology results for 0–3 ungapped mismatches.

**S9 Dataset.** The nine AFR-observed variants that satisfied the limited chromosome 21– restricted, nominal zero-bystander and guide-homology screening criteria.

**S10 Dataset.** The six exploratory AFR-elevated chromosome 21 variants and associated statistical and clinical annotation flags.

**S11 File.** Research analysis scripts, reproducibility checks, data dictionary, provenance notes and SHA-256 manifest.

## References

1. Komor AC, Kim YB, Packer MS, Zuris JA, Liu DR. Programmable editing of a target base in genomic DNA without double-stranded DNA cleavage. Nature. 2016;533:420–424. doi:10.1038/nature17946.

2. Gaudelli NM, Komor AC, Rees HA, et al. Programmable base editing of A•T to G•C in genomic DNA without DNA cleavage. Nature. 2017;551:464–471. doi:10.1038/nature24644.

3. Rees HA, Liu DR. Base editing: precision chemistry on the genome and transcriptome of living cells. Nat Rev Genet. 2018;19:770–788. doi:10.1038/s41576-018-0059-1.

4. Anzalone AV, Randolph PB, Davis JR, et al. Search-and-replace genome editing without double-strand breaks or donor DNA. Nature. 2019;576:149–157. doi:10.1038/s41586-019-1711-4.

5. Landrum MJ, Lee JM, Benson M, et al. ClinVar: improving access to variant interpretations and supporting evidence. Nucleic Acids Res. 2018;46:D1062–D1067. doi:10.1093/nar/gkx1153.

6. gnomAD Production Team. gnomAD v4.1 release announcement. gnomAD Browser. 19 April 2024. https://gnomad.broadinstitute.org/news/2024-04-gnomad-v4-1 (accessed 5 November 2026).

7. Gudmundsson S, Singer-Berk M, Watts NA, et al. Variant interpretation using population databases: lessons from gnomAD. Hum Mutat. 2022;43:1012–1030. doi:10.1002/humu.24309.

8. Fatumo S, Chikowore T, Choudhury A, Ayub M, Martin AR, Kuchenbaecker K. A roadmap to increase diversity in genomic studies. Nat Med. 2022;28:243–250. doi:10.1038/s41591-021-01672-4.

9. Tishkoff SA, Reed FA, Friedlaender FR, et al. The genetic structure and history of Africans and African Americans. Science. 2009;324:1035–1044. doi:10.1126/science.1172257.

10. Richter MF, Zhao KT, Eton E, et al. Phage-assisted evolution of an adenine base editor with improved Cas domain compatibility and activity. Nat Biotechnol. 2020;38:883–891. doi:10.1038/s41587-020-0453-z.

11. Arbab M, Shen MW, Mok B, et al. Determinants of base editing outcomes from target library analysis and machine learning. Cell. 2020;182:463–480.e30. doi:10.1016/j.cell.2020.05.037.

12. Marquart KF, Allam A, Janjuha S, et al. Predicting base editing outcomes with an attention-based deep learning algorithm trained on high-throughput target library screens. Nat Commun. 2021;12:5114. doi:10.1038/s41467-021-25375-z.

13. Morales J, Pujar S, Loveland JE, et al. A joint NCBI and EMBL-EBI transcript set for clinical genomics and research. Nature. 2022;604:310–315. doi:10.1038/s41586-022-04558-8.

14. Bae S, Park J, Kim JS. Cas-OFFinder: a fast and versatile algorithm that searches for potential off-target sites of Cas9 RNA-guided endonucleases. Bioinformatics. 2014;30:1473–1475. doi:10.1093/bioinformatics/btu048.

15. Benjamini Y, Hochberg Y. Controlling the false discovery rate: a practical and powerful approach to multiple testing. J R Stat Soc B. 1995;57:289–300. doi:10.1111/j.2517-6161.1995.tb02031.x.

16. Clopper CJ, Pearson ES. The use of confidence or fiducial limits illustrated in the case of the binomial. Biometrika. 1934;26:404–413. doi:10.1093/biomet/26.4.404.

17. McLaren W, Gil L, Hunt SE, et al. The Ensembl Variant Effect Predictor. Genome Biol. 2016;17:122. doi:10.1186/s13059-016-0974-4.

18. Richards S, Aziz N, Bale S, et al. Standards and guidelines for the interpretation of sequence variants. Genet Med. 2015;17:405–424. doi:10.1038/gim.2015.30.

19. gnomAD Production Team. Genetic ancestry: terminology and classification in gnomAD. 2023. https://gnomad.broadinstitute.org/news/2023-11-genetic-ancestry (accessed 5 November 2026).

20. NCBI ClinVar. Review status documentation. https://www.ncbi.nlm.nih.gov/clinvar/docs/review_status/ (accessed 5 November 2026).

21. UCSC Genome Browser. Human reference assembly hg38 chromosome FASTA. https://hgdownload.soe.ucsc.edu/goldenPath/hg38/chromosomes/chr21.fa.gz (accessed 5 November 2026).

22. Koblan LW, Doman JL, Wilson C, et al. Improving cytidine and adenine base editors by expression optimization and ancestral reconstruction. Nat Biotechnol. 2018;36:843–846. doi:10.1038/nbt.4172.

23. Huang TP, Zhao KT, Miller SM, et al. Circularly permuted and PAM-modified Cas9 variants broaden the targeting scope of base editors. Nat Biotechnol. 2019;37:626–631. doi:10.1038/s41587-019-0134-y.

24. Karczewski KJ, Francioli LC, Tiao G, et al. The mutational constraint spectrum quantified from variation in 141,456 humans. Nature. 2020;581:434–443. doi:10.1038/s41586-020-2308-7.

25. Martin AR, Kanai M, Kamatani Y, et al. Clinical use of current polygenic risk scores may exacerbate health disparities. Nat Genet. 2019;51:584–591. doi:10.1038/s41588-019-0379-x.

26. Choudhury A, Aron S, Botigué LR, et al. High-depth African genomes inform human migration and health. Nature. 2020;586:741–748. doi:10.1038/s41586-020-2859-7.

27. Bergström A, McCarthy SA, Hui R, et al. Insights into human genetic variation and population history from 929 diverse genomes. Science. 2020;367:eaay5012. doi:10.1126/science.aay5012.

28. Musunuru K, Grandinette SA, Wang X, et al. Patient-Specific In Vivo Gene Editing to Treat a Rare Genetic Disease. N Engl J Med. 2025;392:2235–2243. doi:10.1056/NEJMoa2504747.

29. Gupta AO, Sharma A, Frangoul H, et al. Base Editing of HBG1 and HBG2 Promoters for Sickle Cell Disease. N Engl J Med. 2026;394:1824–1835. doi:10.1056/NEJMoa2504835.

30. Frangoul H, Locatelli F, Sharma A, et al. Exagamglogene Autotemcel for Severe Sickle Cell Disease. N Engl J Med. 2024;390:1649–1662. doi:10.1056/NEJMoa2309676.

31. Kim D, Lim K, Kim ST, et al. Genome-wide target specificities of CRISPR RNA-guided programmable deaminases. Nat Biotechnol. 2017;35:475–480. doi:10.1038/nbt.3852.

32. Grünewald J, Zhou R, Iyer S, et al. CRISPR DNA base editors with reduced RNA off-target and self-editing activities. Nat Biotechnol. 2019;37:1041–1048. doi:10.1038/s41587-019-0236-6.

33. Zhou C, Sun Y, Yan R, et al. Off-target RNA mutation induced by DNA base editing and its elimination by mutagenesis. Nature. 2019;571:275–278. doi:10.1038/s41586-019-1314-0.

34. Jaganathan K, Kyriazopoulou Panagiotopoulou S, McRae JF, et al. Predicting Splicing from Primary Sequence with Deep Learning. Cell. 2019;176:535–548.e24. doi:10.1016/j.cell.2018.12.015.

35. Clement K, Rees H, Canver MC, et al. CRISPResso2 provides accurate and rapid genome editing sequence analysis. Nat Biotechnol. 2019;37:224–226. doi:10.1038/s41587-019-0032-3.

36. Tsai SQ, Zheng Z, Nguyen NT, et al. GUIDE-seq enables genome-wide profiling of off-target cleavage by CRISPR-Cas nucleases. Nat Biotechnol. 2015;33:187–197. doi:10.1038/nbt.3117.

37. Scholefield J, Harrison PT. Prime editing—an update on the field. Gene Ther. 2021;28:396–401. doi:10.1038/s41434-021-00263-9.

38. Chen PJ, Liu DR. Prime editing for precise and highly versatile genome manipulation. Nat Rev Genet. 2023;24:161–177. doi:10.1038/s41576-022-00541-1.

39. Gillmore JD, Gane E, Taubel J, et al. CRISPR-Cas9 In Vivo Gene Editing for Transthyretin Amyloidosis. N Engl J Med. 2021;385:493–502. doi:10.1056/NEJMoa2107454.

40. Locatelli F, Lang P, Wall D, et al. Exagamglogene Autotemcel for Transfusion-Dependent β-Thalassemia. N Engl J Med. 2024;390:1663–1676. doi:10.1056/NEJMoa2309673.

41. Hu JH, Miller SM, Geurts MH, et al. Evolved Cas9 variants with broad PAM compatibility and high DNA specificity. Nature. 2018;556:57–63. doi:10.1038/nature26155.

